# Identifying robust functional modules using three-body correlations in *Escherichia coli*

**DOI:** 10.1101/2021.08.03.454257

**Authors:** Tianlong Chen, Pramesh Singh, Kevin E. Bassler

## Abstract

Understanding the underlying structure of a gene regulatory network is crucial to understand the biological functions of genes or groups of genes. A common strategy to investigate it is to find community structure of these networks. However, methods of finding these communities are often sensitive to noise in the gene expression data and the inherent stochasticity of the community detection algorithms. Here we introduce an approach for identifying functional groups and their hierarchical organization in gene co-expression networks from expression data. A network describing the relatedness in the expression profiles of genes is first inferred using an information theoretic approach. Community structure within the inferred network is found by using *modularity maximization*. This community structure is further refined using three-body structural correlations to robustly identify important functional gene communities. We apply this approach to the expression data of *Escherichia coli* genes and identify 25 robust groups, many of which show key associations with important biological functions as demonstrated by Gene Ontology (GO) term enrichment analysis. Thus, our approach makes specific and novel predictions about the function of these genes.

## 1. Introduction

Recent advancements in application of high-throughput techniques such as sequencing and gene expression profiling have resulted in abundance of genetic data. A variety of computational methods and tools have been developed for analyzing these data [1–5]. Robust techniques have been developed to identify the functional and regulatory regions (genes) of sequenced genomes in an automated way [6–9]. However, determining how genes interact and operate together to control the function and behavior at the system level still remains a challenging task. Developing computational methods that use gene expression data (microarray or RNA-Seq data) to identify co-expressed gene modules or communities that act as functional units to predict genes functions and their interactions with other genes is an important step in attaining a systems level understanding of cellular processes. In this paper, we present an information theoretic, network based approach that we apply to investigate the network of gene interactions in *Escherichia coli* (*E. coli*). We identify 25 robust gene communities that significantly enrich the Gene Ontology (GO) terms. These communities include many unannotated genes. The knowledge of their community membership can help us make specific predictions about their function.

Network based approaches to analyze gene expression data have received considerable interest in the field of systems biology [10]. In recent years, a number of network inference (NI) methods have been developed to construct networks that quantify the relationship of genes in the data. Most gene expression data are static data that are the average levels of expression under certain steady conditions. With this type of data, undirected networks with links describing the relatedness in the expression of each pair of genes can be constructed. Methods to construct relatedness networks have been devised that use a variety of similarity measures in gene expression [11–19]. A typical implementation of these methods constructs a weighted network of genes by performing a pairwise comparison of gene expression profiles of all genes. Pairs of genes are connected by links whose weights are determined by the corresponding similarity measure. The Context Likelihood of Relatedness (CLR) is a commonly used method for this purpose [12]. It considers the expression profile of genes in a series of experiments performed under conditions, such as different temperatures, sugars, salinity, knock-outs, or the presence of antibiotics, as a stochastic signal. The first step is to compute an information-theoretic measure called mutual information (MI) [20], which quantifies the correlations in the expression patterns of genes and is a commonly used similarity measure. After that, a “relatedness” score for each pair of genes is calculated from its MI relative to the distribution of MI for each gene in the pair. CLR has been shown to more clearly identify significant relationships than simply using a linear correlation or mutual information [12]. The estimates of gene expression levels in experimental data are inherently noisy and therefore the link weights in the inferred network also have some level of noise. An effective strategy to deal with this problem is to filter out the low weight links in the network inferred using CLR or similar methods by using an appropriate threshold [21].

The expression profiles of genes that share functional similarity is highly correlated. Conversely, similarity between genes (the inferred gene co-expression network) can be used to identify these functional groups. This is where community detection algorithms turn out to be useful as they can extract these communities from a given network. Many methods of community detection exist [22], each of which finds a different type of community [23–28]. We use an unsupervised and intuitively appealing approach to find communities that are more densely connected than at random. This is done by searching for the partition of genes that maximizes a metric called modularity [22,29,30]. Communities of genes that maximize the modularity of the relatedness network have been shown to be effective predictors of function [21]. However, the algorithmic detection of community structure assigns every gene to some community with varying degree of confidence. Sometimes these methods also find too many communities, not all of which are biologically meaningful. In practice, with too many predicted functional associations, it becomes difficult to assess which of these predictions are most reliable, or most important. In fact, many detected gene communities whose structures are statistically significant may not be so functionally. To overcome this practical challenge, it is desirable to have a method that finds a fewer but highly robust gene communities that can be tested for predictions by eliminating genes that are weakly associated.

In this paper, we introduce a method to identify highly robust communities within the network, which we apply to investigate the structure of the gene co-expression network in *E. coli*. We infer the weighted relatedness network using the CLR method, and then analyze it with a modularity maximizing community detection method to find the hierarchical organization of gene communities. Due to the inherent noise in the expression data and the stochastic nature of community detection algorithms, the detected communities are slightly noisy. We first use three-body correlations in the community structure to filter the structural information, revealing a set of strong communities. The structure of each of these communities are then individually analyzed by removing weaker links. In doing so, most of the communities fall apart, but we find small set of 25 core communities that remain largely intact. Focusing on these highly stable communities, we perform a *Gene Ontology (GO) term enrichment analysis* for their functional annotations. Further, we also analyze the known annotations of the genes in the communities, allowing us to make novel predictions for the function and interactions of specific genes. Although our analysis focuses on the model organism *E. coli*, the approach presented here for analyzing gene co-expression networks is more general and can be applied to study other types of molecular interaction networks as well as other organisms.

## 2. Methods

### 2.1. Data

We analyze version 4, build 6 gene expression data from the Many Microbe Microarrays Database(*M* ^3*D*^) [31]. This dataset is a compendium of expression profiles for 4297 *E. coli* MG1655 genes resulting from 907 experiments. These experiments were conducted under 466 different conditions, such as temperatures, pH levels, presence of antibiotics, oxygen concentration, genetic perturbations etc. The expression levels derived from different experimental conditions have been normalized for each gene.

To perform the enrichment analysis, we use the Gene Ontology data available from Gene Ontology Consortium [32]. We also compare our clustering results against the RegulonDB data, which is the primary database of the transcriptional regulation in *E. coli* K-12. The data and knowledge in both of these databases is manually curated from original scientific publications.

### 2.2. Network inference

From the expression profiles of genes we infer the gene regulatory networks using the context likelihood of relatedness (CLR) method. The CLR algorithm is an extension of the traditional relevance network method [12]. Like relevance networks, CLR algorithm first calculates a metric of similarity between the expression profiles of each pair of genes. The metric chosen here is the mutual information (MI). Mutual information is shown [12] to be a better measure than common statistical metrics like the Pearson correlation, as it takes into account pairwise nonlinear relationships between variables. The mutual information of the expression profiles of two genes is calculated as:

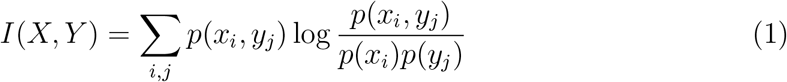

where *p*(*x_i_*) is the marginal probability that *X* = *x_i_* and *p*(*x_i_, y_j_*) is the joint probability that *X* = *x_i_, Y* = *y_j_*. To get these probability values from the original expression profiles, a B-spline smoothing and discretization method was applied [20]. We used the open uniform knot vector with the number of bins = 10 and the spline order = 3. For *N* genes, we obtain *I* as a symmetric *N* × *N* matrix.

Indirect regulatory influences and direct physical interactions between gene pairs are not easy to distinguish due to bias in the expression profiles data. Such bias normally come from uneven condition sampling, upstream regulation and inter-laboratory variations in microarrays. The CLR algorithm tries to solve this problem by going a further step that takes the network context of each relationship into account. It compares the similarity metric of one pair of genes, i and j, to its corresponding background distribution. Thus, MI between gene pair X and Y *I*(*X, Y*) is compared to the distribution of MI between gene X(or Y) and all other genes {*I*(*X, Y*); ∀*Y* (*or* ∀*X*)}. A z-score is then calculated using:

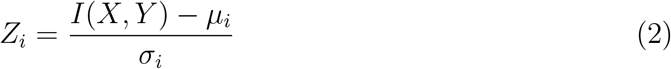

where *μ_i_* and *σ_i_* are the mean and standard deviation of the corresponding distribution. Negative z-scores are set to be zero. Then the final likelihood value is defined as:

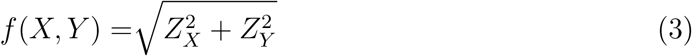

The resultant symmetric matrix of *f* (*X, Y*) is called the relatedness matrix and is regarded as the weighted adjacency matrix for the regulatory network (with scores along the diagonal set to zero). This further step has been proved to have increased the contrast between the physical interactions and the indirect relationship, which reduces the noise in the regulatory network inferred.

### 2.3. Community detection

We use modularity to study the community structure of the inferred regulatory network. Modularity is a widely used measure to quantify the strength of division of a network into modules [30]. Mathematically, modularity *Q* is defined as

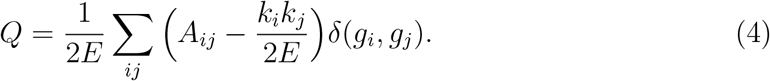

here *A* is the adjacency matrix, *k_i_*,*k_j_* is the degree of node *i* and *j*, *m* is the total number of links in the network, i.e. 2*E* = Σ*_i_ k_i_*, *δ* is the Kronecker delta function and *g_i_* and *g_j_* refer to the communities that nodes *i* and *j* belong to. For weighted networks, *Q* is defined in the same way as above except *A_ij_* is the element of weighted adjacency matrix, *k_i_* and *k_j_* are the sums of the link weights attached to nodes *i* and *j* respectively, and *E* is the sum of weights of all links in the network.

For a given network, the partition that has the highest value of the modularity *Q*_max_ determines how modular the network is. The goal of a community detection method is to find the partition of network nodes which has the highest modularity, which represents the communities of the network. Unfortunately, this has been proven to be an NP-hard problem [33]. In recent years, a number of heuristic algorithms have been developed to find a network partition with modularity close to its maximum value [29, 30, 34–36]. A recent method which combines the leading eigenvalue method of [30] with final tuning has been proved to eliminate detection bias [37] and is shown to performs well in detecting communities [21]. Here we applied a variant of this algorithm with an extra agglomeration step described in [38] which show significantly improved results when tested on a variety of benchmark networks.

### 2.4. Co-clustering analysis

The relatedness matrix represents a weighted network which is dense and contains a large number of low-weight links. We discard (set their weights to 0) the links whose relatedness value is smaller than a chosen threshold *f*_min_. In this way, a series of relatively sparse weighted networks were derived from the original single weighted network using a different *f*_min_.

The community detection algorithm described above is then applied to study the series of networks. Due to the stochastic nature of the algorithm, which is common feature of most community detection methods, the partitions found are different for different runs. To account for this variability, we conduct a statistical analysis based on the ensemble of partitions obtained from running the algorithm 1000 times. Specifically, a ‘correlation’ value of each pair of genes is defined as the chance they were discovered in the same community. For each *f*_min_ value, we obtain a *N* × *N* symmetric co-cluster matrix representing the co-occurrence probability for each pair of genes, where *N* is the number of genes in the network. These matrices can be visualized as a color map in the in the following way. Since we have three different co-cluster matrices (for three different *f*_min_), we could use a different color (Red, Green, or Blue) to represent each of them. For each co-cluster matrix, the strength of the color proportional to the co-clustering probability corresponding to the matrix element. In this way, the similarity between a pair of genes in three different networks can be combined to a form a pixel color and all three co-cluster matrices can be visualized as a single 2-dimensional color map.

Here we focus on the three networks corresponding to *f*_min_ = 3, 4 and 5 (sub-critical, critical, and super-critical thresholds). The critical value (below which the size of the largest connected component is the size of the network) with respect to the connectivity of the whole network is at about *f* = 4 (Fig. 1). We assign blue (B), red (R), and green (G) for *f*_min_ = 3, 4 and 5 respectively. Each pixel vector containing three values in B, R, and G dimensions allows us to visualize the information of the gene pair’s co-cluster correlation (henceforth referred to simply as correlation) in three different networks simultaneously. A color map (see Fig. 2) is used to visualize the matrix consisting of all pixel vectors for every pair of genes [21].

**Figure 1:**
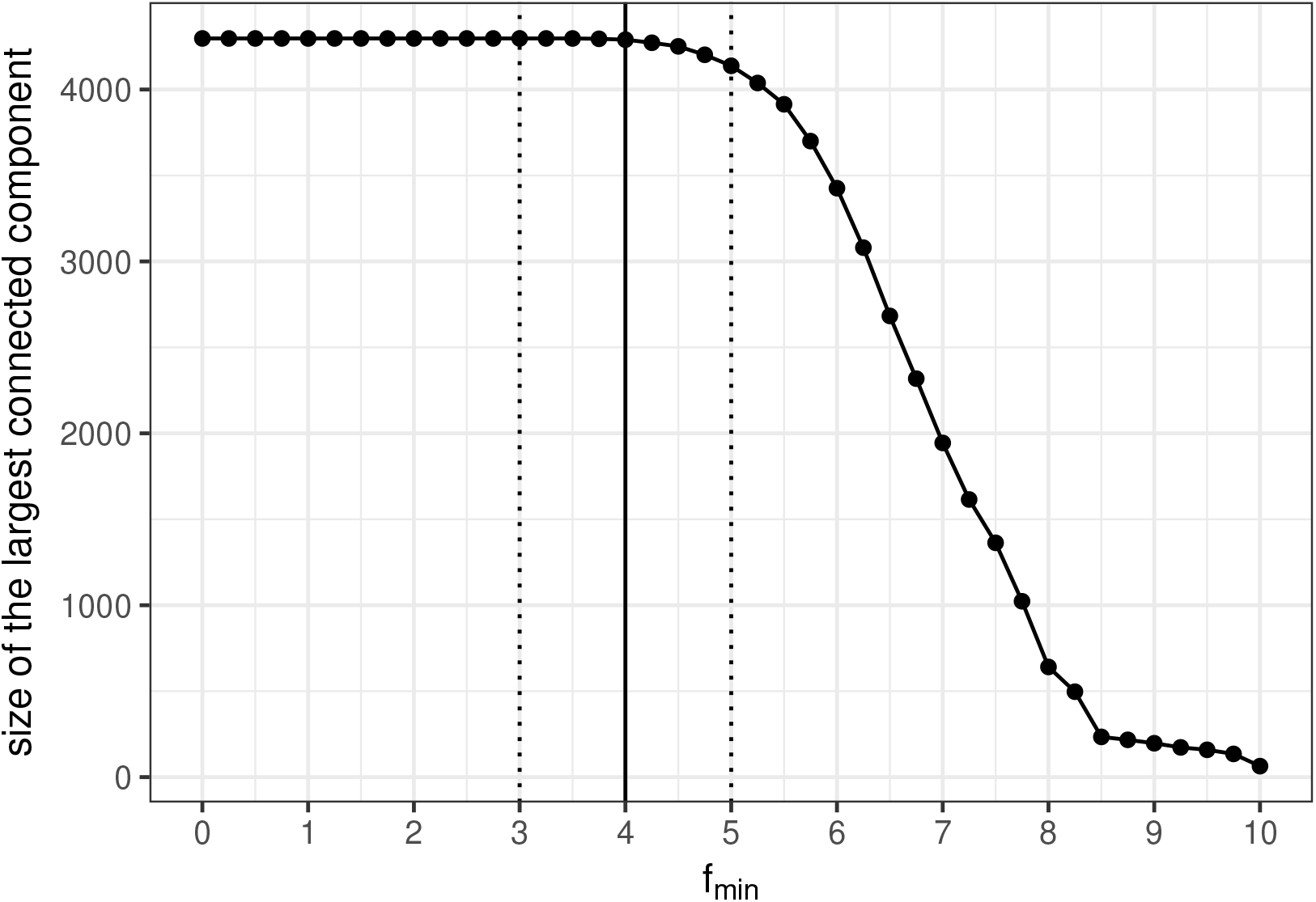
Network connectivity as a function of relatedness threshold. The size of the largest connected component in the network as a function of *f*_min_. As *f*_min_ increases, more links are removed and around *f*_min_ = 4, the network begins to fall apart.

**Figure 2:**
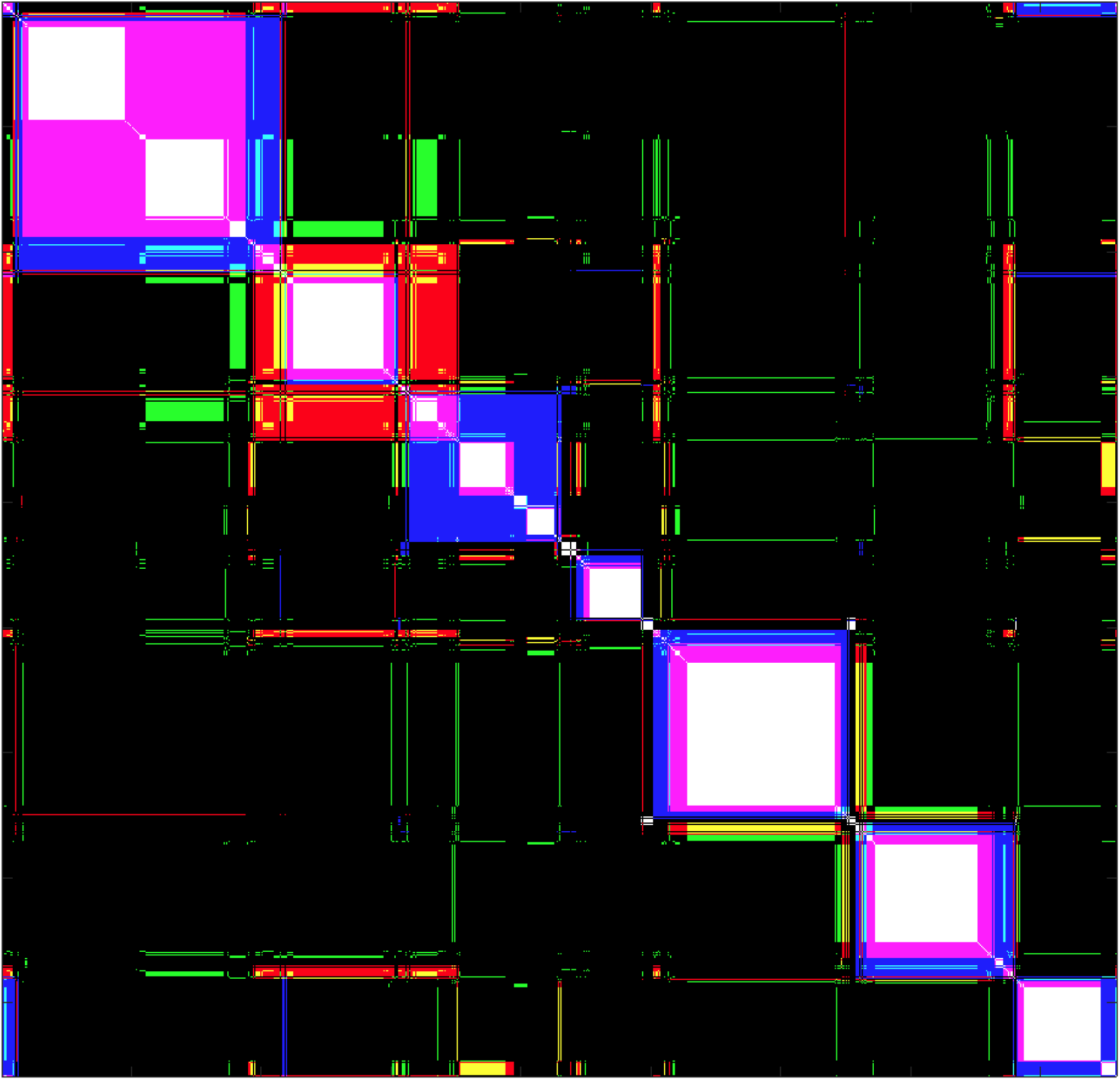
Visualization of co-clustering matrices. The figure is combining three 4297 × 4297 co-cluster matrices resulting from community detection on the networks corresponding to *f*_min_ = 3, 4 and 5. Each pixel at (*i, j*) in the matrix represents co-cluster correlation of the *i*-th and *j*-th genes. If a pixel is blue/red/green, then the corresponding gene pair is found together in the same community at *f*_min_ = 3/4/5. When a gene pair is together in multiple networks, the pixel’s color appears as the superposition of different colors weighted by their co-occurrence frequency. When a gene pair is present in all three communities, the resulting color is white.

### 2.5. Three-body correlation analysis

A typical way to help identify useful patterns from in a network is to reduce noise by eliminating low-information links. For the three co-cluster matrices, we a cut-off close to 1 (we choose 0.75) to be the criterion of the correlation values, for each *f*_min_. When a pair of genes’ correlation exceeds this criterion, they are considered as closely connected in the network for that particular *f*_min_.

Instead of looking at the 4297 genes’ correlations all at once, we take a different approach and view the gene relationships from each individual gene’s perspective. By focusing on one particular gene at a time in the network, we are able to further utilize the correlation information with a three-body analysis. As has been discussed, a pixel (*i, j*) in the co-cluster matrix describes the correlation of gene *i* and *j*. Hence it is a two-body correlation, but we can extend this idea and focus on an objective gene *X* consider the correlation information of gene *i* and *j* only when *X* is connected to them, which then becomes a three-body correlation. Figure 3 illustrates how the looking at the three-body correlations can be helpful to eliminate unimportant information and provide further insight to the underlying network structure. Fig. 3 (a) shows the two-body correlation among genes which are closely connected with our objective gene (*lexA*). Fig. 3(b) shows a three-body correlation map when the pair of gene of each pixel plus the objective gene are found together. Fig. 3(b) is similar to Fig. 3(a), except the regions of correlations not involving the objective gene are removed.

**Figure 3:**
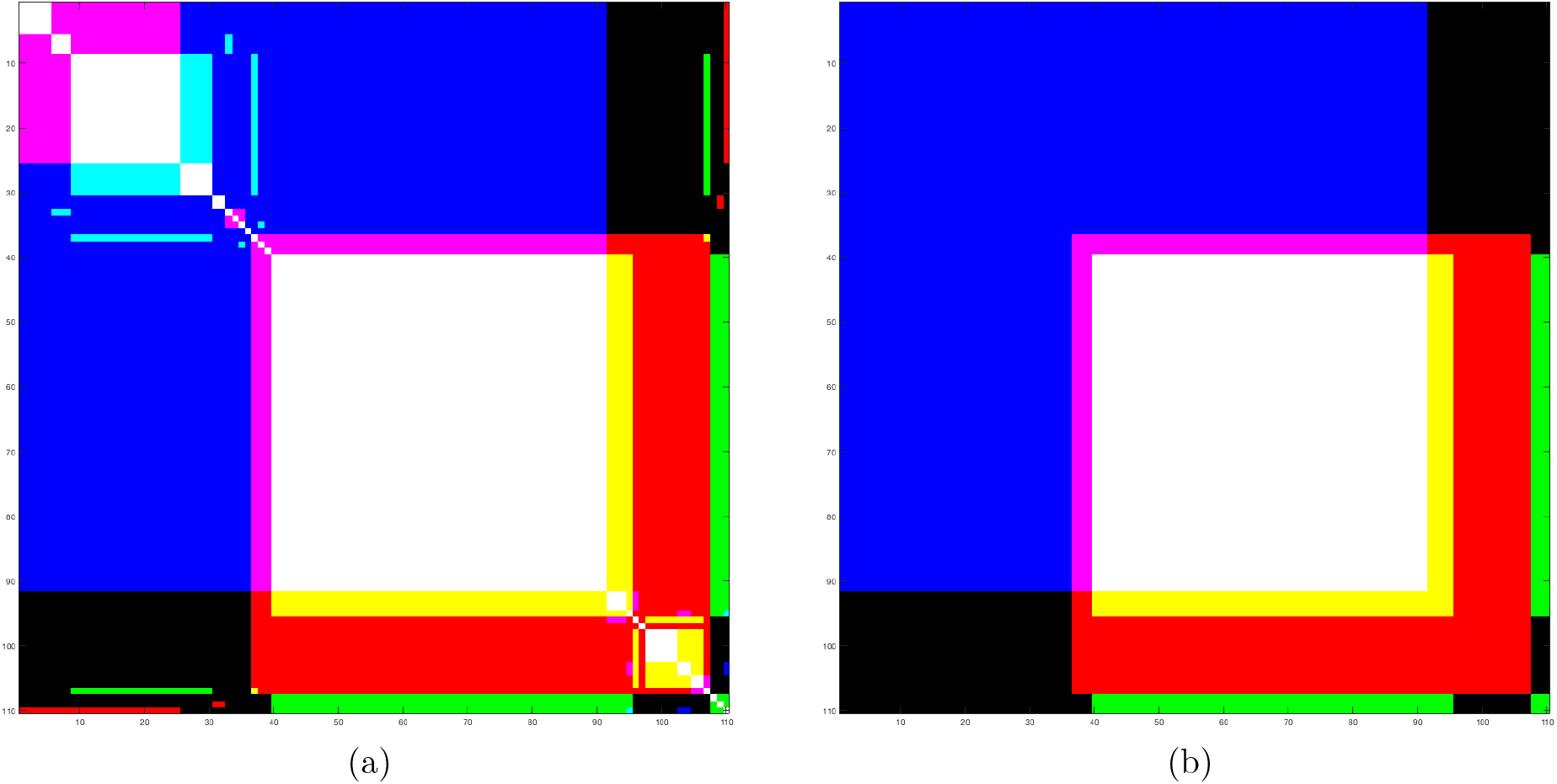
Two-body and three-body co-cluster matrices of 110 genes closely connected with *lexA*. Genes that are closely connected to *lexA* in at least one of the three *f*_min_ networks. These genes are ordered in the same sequence from along the X and Y axes from the upper left corner. This order is kept the same for all *f*_min_ values. The element would be colored using blue, red and green When its correlation value exceeds 0.75 at *f*_min_ =3, 4 and 5 respectively. Here purple (yellow) is blue plus red (green plus red), which means the element was colored at thresholds 3 and 4 (4 and 5). White represents a mix of all three colors (at all three thresholds), while black implies an absence of the three colors. (a) Each pixel (*i, j*) represents the co-cluster correlation between genes *i* and *j* (two-body correlation). (b) Each pixel (*i, j*) here denotes the probability that gene *i*, *j* and *lexA* are found in the same community (three-body correlation).

Using three-body correlation matrix instead of the ordinary two-body correlation is advantageous as it eliminates the information between pairs of non-objective genes and can show clearer structures on the color map. In Fig. 3(b), we find three blocks representing the results from networks with *f*_min_ equals to 3, 4 and 5. The existence of a ‘white block’ can clearly be found in the central area. A white block is formed by high correlation values (normally set between 0.75 and 1.0) in networks at all three *f*_min_ (corresponding to red, green and blue). We particularly focus on the objective genes that show such a white block since it implies presence of a robust group of genes.

### 2.6. Simulated annealing to visualize co-cluster matrices

Simulated annealing is a widely used method for finding an approximate solution to non-linear optimization problems [39]. Here we use it to find optimal ordering of genes to visualize the gene groups in a blockdiagonal form on a 2-dimensional color map. The objective function that we use is

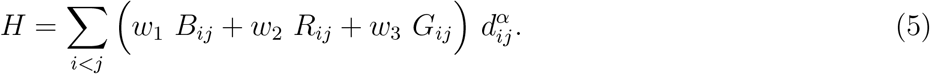

The element (*i, j*) of a co-cluster matrix measures the fraction of times gene *i* and gene *j* are found in the same community in an ensemble of community detection results obtained by a stochastic algorithm like the one used in this paper. *B_ij_* corresponds to the smallest threshold value and determines the strength of color blue when mapped visually. Similarly *R_ij_* (red) and *G_ij_* (green) corresponds to middle and the largest threshold values respectively. *d* is a matrix measuring distance of an element (*i, j*) from the matrix diagonal (*i* = *j*).

Starting from a randomly ordered set of genes (while maintaining the same order along the row and the column) in the co-cluster matrix, we use the Metropolis algorithm [39] to minimize the function *H*. The algorithm starts from an initial temperature *T* (chosen to be very high) and at each step a the order of an adjacent pair of genes is interchanged. If this step results in a decreased value of *H* then it is accepted, otherwise the change Δ*H* is accepted with a probability proportional to *e*^−Δ*H/T*^ while the temperature is gradually lowered according to a prescribed cooling schedule. Our choice of *α* = 1.0, *w*_1_ = 0.25, *w*_2_ = 0.5, *w*_3_ = 1.0, with a varying cooling schedule gives a satisfactory convergence and a reasonable approximation to the optimal ordering.

### 2.7. Further pruning gene groups

An increase in *f*_min_ results in disappearance of a number of links in the network. With increasing *f*_min_, we expect the network to reach a stage where it no longer has a giant component (a connected component with size of the order of network size *N*) and the network begins to fragment into smaller disconnected components. For the *E. coli* relatedness network, this happens around *f*_min_ = 6. At higher thresholds, it is simpler and efficient to look at the links among genes within the white block instead of using community detection algorithms. Thus, we focus our attention to the connected components within the white blocks. By increasing *f*_min_ and thereby removing links between genes, we observe how the connected components gradually fragments, and identify the most robust group containing the objective gene.

Finally, we conduct a genome wide study by systematically treating every gene in the *E. coli* network as the objective gene and repeating the procedure outlined above. It turned out that our approach could not guarantee to discover a possible functional unit for every gene. For some genes, either a clear white block was not found or their connected components within the white block became disconnected at relatively smaller values of *f*_min_. However, our approach works well in a number of objective genes. We identified 25 gene groups that contained a large fraction of their genes of their corresponding white blocks at a high threshold of *f*_min_ = 8 (Supplementary Material S1). Most of these groups provide significant GO term enrichment.

### 2.8. GO term enrichment analysis

We compare the detected gene communities with known Gene Ontology (GO) terms [32] using the hypergeometric test. Each community is compared with a GO term and then the p-value for overrepresentation is computed as

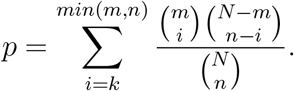

Here *n* is the number of genes in the community, *m* is the number of genes in the GO term, *k* is the number of genes common between the community and the GO term, and *N* is the total population (intersection set) of genes present in both sets (in the GO term data as well as in the communities). For a fast and efficient computation of the p-values we used a java based tool *Ontologizer* [40]. Since we are performing multiple tests, we further adjust these p-values by applying the Benjamini-Hochberg correction [41].

## 3. Results

### 3.1. Genome wide analysis

By systematically treating all the genes as objective genes, we identify 25 stable potential functional units. To obtain these groups we make a slight variation in the community detection algorithm [38], which yields better results (more significant enrichment and more robust communities). In the bisection step of the algorithm, instead of performing the division according to the leading eigenvalue we divide the groups randomly which results in more significant and stable groups. These communities also contain stable subgroups of genes that remains connected at extremely high values of *f*_min_. These identified groups have been found to be associated (observed in terms of significant number of common genes) with certain biological functions as listed in Table 1. We name some of these groups according to their known function.

**Table 1:** The 25 identified robust groups labeled by their known biological functions along with the size of the groups. The stable component is the size of the connected component within white block at very high thresholds (*f*_min_ = 8).

| Group | White-block size | Stable component |
| --- | --- | --- |
| Unknown | 250 | 112 |
| Amino acid | 200 | 144 |
| Alternative carbon | 184 | 110 |
| Anaerobic | 100 | 86 |
| Mobility | 70 | 67 |
| LexA | 50 | 44 |
| Carbon starvation | 41 | 32 |
| Iron deficiency | 47 | 43 |
| Heat shock | 27 | 16 |
| Unknown | 17 | 14 |
| tRNAs | 15 | 14 |
| Unknown | 15 | 8 |
| Unknown | 10 | 10 |
| Cytochrome bo oxidase complex | 8 | 5 |
| ATP uptake | 10 | 7 |
| Unknown | 8 | 8 |
| Fe S cluster biosynthesis | 7 | 6 |
| Unknown | 7 | 7 |
| Unknown | 7 | 6 |
| Unknown | 7 | 7 |
| Unknown | 6 | 6 |
| Unknown | 5 | 5 |
| Unknown | 5 | 5 |
| Phosphate uptake | 5 | 5 |

To test the biological significance of the white blocks and the further identified 25 gene groups, we conduct enrichment analysis and look at the statistical overrepresentation of the gene clusters with *Gene Ontology* (GO) database. We also compare these groups with the latest experimental datasets of *Operon* and regulatory network data (*Transcription Factor (TF)-Operon* network) from the *RegulonDB* database (version 9.4) [42]. An operon is a functional unit of DNA containing one or a few genes under the control of a single promoter. Genes belonging to the same operon are *co-transcribed* and their expression profile are highly correlated. We test the efficiency of our network analysis method by calculating the proportion of completely retrieved operons from the white blocks. The total number of operons, containing at least 2 genes, found in the network is 727. We recover 339 of them (which is about 46.7%); each of which from the same group is in the same white block.

### 3.2. GO term enrichment

In this section we show that the detected communities are biologically relevant by comparing these communities with Gene Ontology (GO) terms. This is done by performing hypergeometric test between gene communities and GO terms. The statistical significance of the matching is indicated by the *p*-value of hypergeometric test (smaller *p*-values indicating higher significant matches). The top few enrichment results are summarized in Table 2 and the most significant enrichment for each group is shown in Table 3. Most significant associations are also visualized in Fig. 4. The same community detection method can be applied to each of these communities separately to detect functional structures within them. A full list of statistically significant associations is included in the Supplemental Material S2.

**Table 2:** Most significant GO term enrichment results. 30 most significant associations between groups and GO terms. m is the size of the GO term, n is the size of the group, k is the size of the overlap, and p is the Benjamini-Hochberg corrected p-value of the hypergeometric test.

| Group | GO term | GO size | Com size | In common | P value |
| --- | --- | --- | --- | --- | --- |
| Amino acid | cellular amino acid biosynthetic process | 105 | 144 | 53 | 1.21E-49 |
| Amino acid | alpha-amino acid biosynthetic process | 91 | 144 | 49 | 1.54E-47 |
| Amino acid | alpha-amino acid metabolic process | 152 | 144 | 57 | 3.43E-45 |
| Amino acid | cellular amino acid metabolic process | 194 | 144 | 62 | 5.44E-45 |
| Amino acid | small molecule biosynthetic process | 226 | 144 | 64 | 4.63E-43 |
| Amino acid | carboxylic acid biosynthetic process | 161 | 144 | 56 | 1.77E-42 |
| Amino acid | organic acid biosynthetic process | 161 | 144 | 56 | 1.77E-42 |
| Mobility | locomotion | 64 | 67 | 30 | 3.26E-37 |
| Amino acid | single-organism biosynthetic process | 415 | 144 | 70 | 7.23E-32 |
| Amino acid | organonitrogen compound biosynthetic process | 394 | 144 | 68 | 2.11E-31 |
| Amino acid | carboxylic acid metabolic process | 394 | 144 | 67 | 1.99E-30 |
| Amino acid | oxoacid metabolic process | 401 | 144 | 67 | 5.78E-30 |
| Amino acid | organic acid metabolic process | 404 | 144 | 67 | 8.68E-30 |
| Amino acid | small molecule metabolic process | 639 | 144 | 81 | 3.55E-29 |
| Anaerobic | anaerobic respiration | 54 | 86 | 25 | 4.09E-27 |
| LexA | SOS response | 22 | 44 | 16 | 2.11E-26 |
| Mobility | localization of cell | 49 | 67 | 22 | 5.18E-26 |
| Mobility | cell motility | 49 | 67 | 22 | 5.18E-26 |
| Mobility | cilium or flagellum-dependent cell motility | 42 | 67 | 21 | 5.18E-26 |
| Mobility | bacterial-type flagellum-dependent cell motility | 42 | 67 | 21 | 5.18E-26 |
| Mobility | archaeal or bacterial-type flagellum-dependent cell motility | 42 | 67 | 21 | 5.18E-26 |
| Anaerobic | energy derivation by oxidation of organic compounds | 97 | 86 | 28 | 5.07E-24 |
| Mobility | movement of cell or subcellular component | 59 | 67 | 22 | 7.24E-24 |
| Anaerobic | generation of precursor metabolites and energy | 113 | 86 | 29 | 2.17E-23 |
| Heat shock | response to heat | 56 | 16 | 14 | 2.35E-23 |
| Anaerobic | cellular respiration | 83 | 86 | 26 | 3.31E-23 |
| Amino acid | organonitrogen compound metabolic process | 761 | 144 | 78 | 1.85E-21 |
| Heat shock | response to temperature stimulus | 76 | 16 | 14 | 2.47E-21 |
| Amino acid | single-organism process | 1710 | 144 | 115 | 5.17E-21 |
| Iron deficiency | siderophore transport | 15 | 43 | 12 | 1.97E-20 |

**Table 3:** GO term enrichment for each group. Most significant association found for each group. m is the size of the GO term, n is the size of the group, k is the size of the overlap, and p is the Benjamini-Hochberg corrected p-value of the hypergeometric test.

| Group | GO term | GO size | Com size | In common | P value |
| --- | --- | --- | --- | --- | --- |
| Unknown | response to stimulus | 781 | 112 | 43 | 5.99E-06 |
| Amino acid | cellular amino acid biosynthetic process | 105 | 144 | 53 | 1.21E-49 |
| Alternative carbon | aerobic respiration | 37 | 110 | 12 | 2.64E-09 |
| Anaerobic | anaerobic respiration | 54 | 86 | 25 | 4.09E-27 |
| Mobility | locomotion | 64 | 67 | 30 | 3.26E-37 |
| LexA | SOS response | 22 | 44 | 16 | 2.11E-26 |
| Iron deficiency | siderophore transport | 15 | 43 | 12 | 1.97E-20 |
| Carbon starvation | ligase activity, forming carbon-sulfur bonds | 25 | 31 | 5 | 1.29E-05 |
| Heat shock | response to heat | 56 | 16 | 14 | 2.35E-23 |
| Unknown | monosaccharide catabolic process | 52 | 14 | 7 | 4.31E-09 |
| Unknown | diaminopropionate ammonia-lyase activity | 1 | 8 | 1 | 1.10E-2 |
| Unknown | [Ni-Fe] hydrogenase complex | 7 | 10 | 3 | 6.55E-06 |
| ATP uptake | heme transport | 9 | 7 | 5 | 1.92E-11 |
| Unknown | ferredoxin hydrogenase complex | 6 | 8 | 5 | 2.60E-12 |
| Cytochrome bo oxidase complex | cytochrome o ubiquinol oxidase complex | 4 | 5 | 4 | 2.89E-11 |
| Unknown | protein-phosphocysteine-galactitol-phosphotransferase system transporter activity | 4 | 7 | 3 | 3.53E-07 |
| Unknown | galactarate metabolic process | 4 | 6 | 4 | 8.28E-11 |
| Unknown | metallo-sulfur cluster assembly | 18 | 7 | 6 | 1.51E-12 |
| Fe S cluster biosynthesis | iron-sulfur cluster transfer complex | 2 | 6 | 2 | 2.77E-05 |
| Unknown | carbohydrate derivative catabolic process | 45 | 6 | 5 | 2.86E-08 |
| Unknown | threonine metabolic process | 12 | 5 | 4 | 8.87E-09 |
| Unknown | (4S)-4-hydroxy-5-phosphonooxypentane-2,3-dione isomerase activity | 1 | 5 | 1 | 7.467E-3 |
| Phosphate uptake | regulation of phosphorus metabolic process | 10 | 5 | 5 | 1.98E-12 |

**Figure 4:**
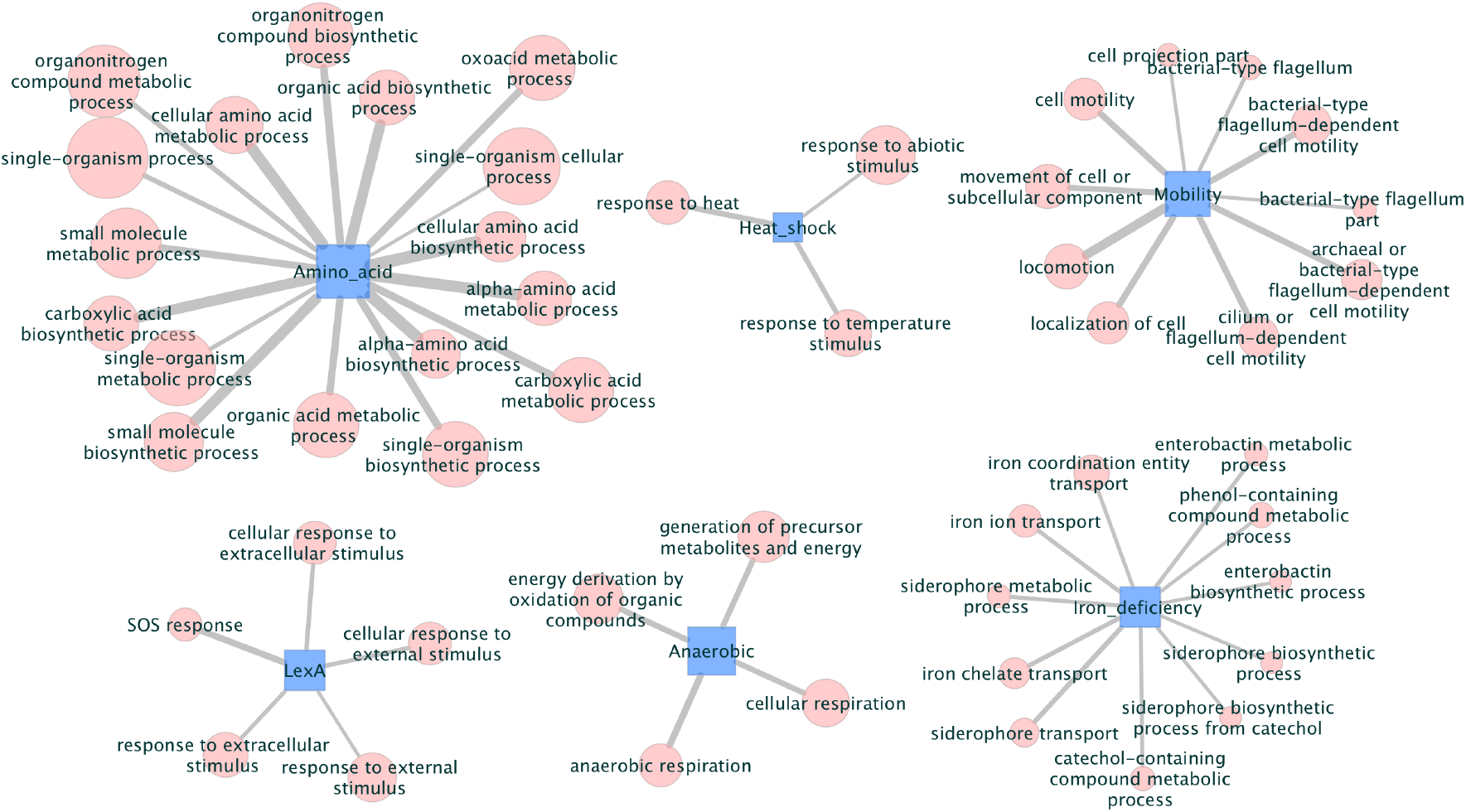
Most significant GO term enrichment. Bipartite network showing the 50 most significant associations between clusters discovered and GO terms. Nodes in blue squares are the clusters and pink circles depict the GO term. The thickness of the links is proportional to the negative logarithm of the p-values obtained by hypergeometric test.

### 3.3. Potential regulatory relationships

We further investigate the 25 gene groups which are identified by increasing *f*_min_ inside the white blocks. We use to the experimental regulatory network data of TFs and operons obtained from RegulonDB. Here we present three example cases, which are described as the ‘Iron deficiency’, ‘LexA’, and ‘Mobility’ groups as shown in Fig. 5, Fig. 6, and Fig. 7 respectively. Most of the genes/operons in these groups are regulated by known TFs. For example in the Iron deficiency group, the TF Fur is associated with siderophore transport [43], and is also revealed by our GO term enrichment analysis (Table 3). Similarly, the transcription factors *LexA* and *FlhDC* are known to regulate genes that are related to SOS response [44] and mobility [45] in *E. coli* respectively. In all three figures(Fig. 5, Fig. 6, and Fig. 7), genes belonging to the same operon are collapsed into the same node. Each pink or green node represents one operon, and their connections based on our original inferred relatedness network are represented by light gray lines. The small blue nodes are all the involved TFs which have regulatory relationship according to the RegulonDB data. Each regulatory relationship is shown using a dark black line. It is reasonable to group these operons (genes) together as they are regulated by the same set of TFs. In addition, we also discover a few nodes (colored green) in our groups which have no connection to any known TFs in the RegulonDB data. Many of these genes have names starting with character ‘y’ and it turns out that the current understanding of these ‘y genes’ is quite limited [46]. Regarding these genes, we assert that our results are indicative of new regulatory relationships and gene functions among the members of these groups.

**Figure 5:**
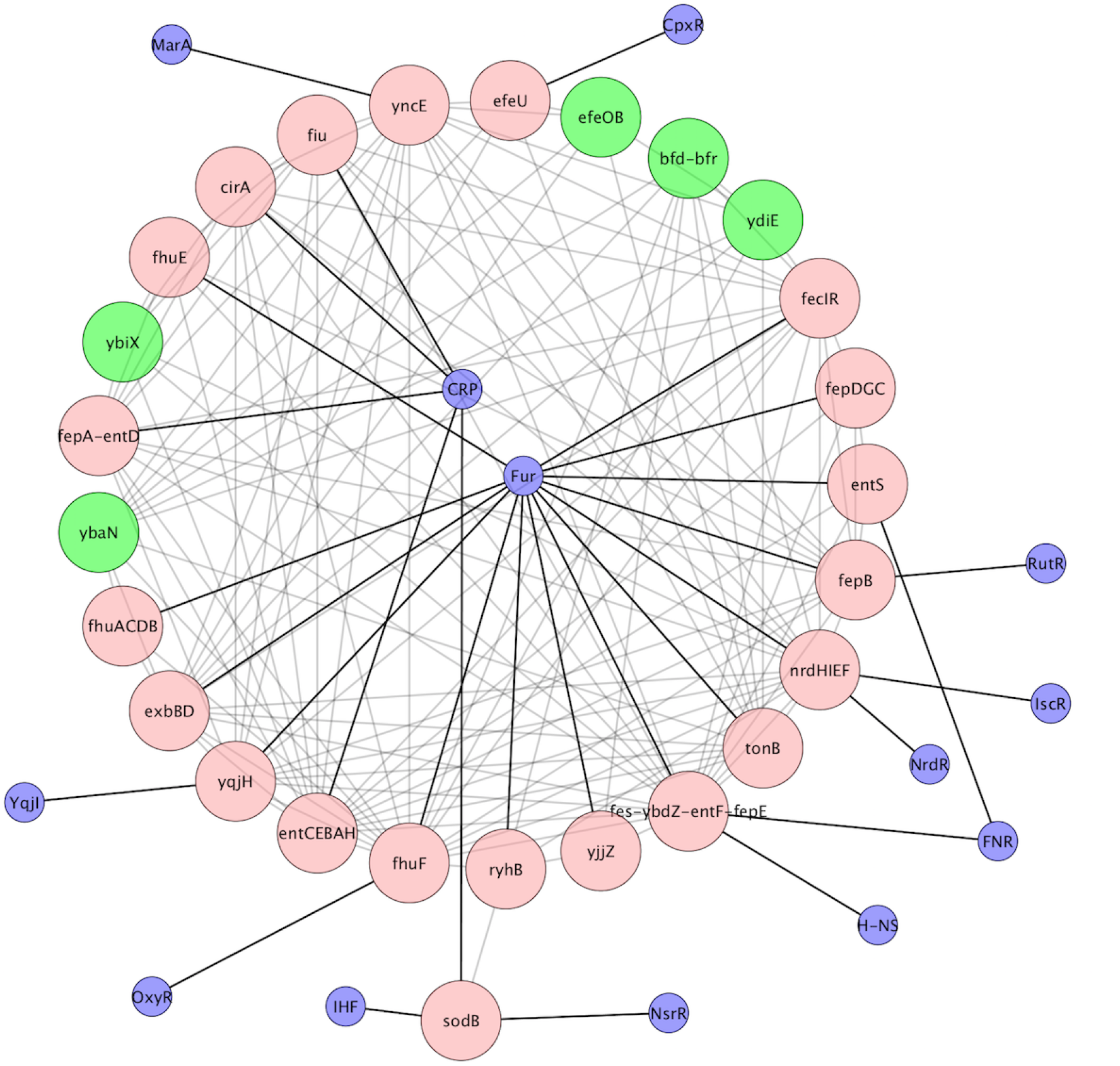
The sub-network of the group ‘Iron deficiency’. The group contains 43 genes. Genes belonging to the same operon are collapsed into the same node in the figure. Pink and green nodes are the members in our group. The small blue nodes are TFs which have regulatory connection with the operons (genes). Links between genes are shown as light gray lines and the regulatory relationship from RegulonDB is drawn as black lines.

**Figure 6:**
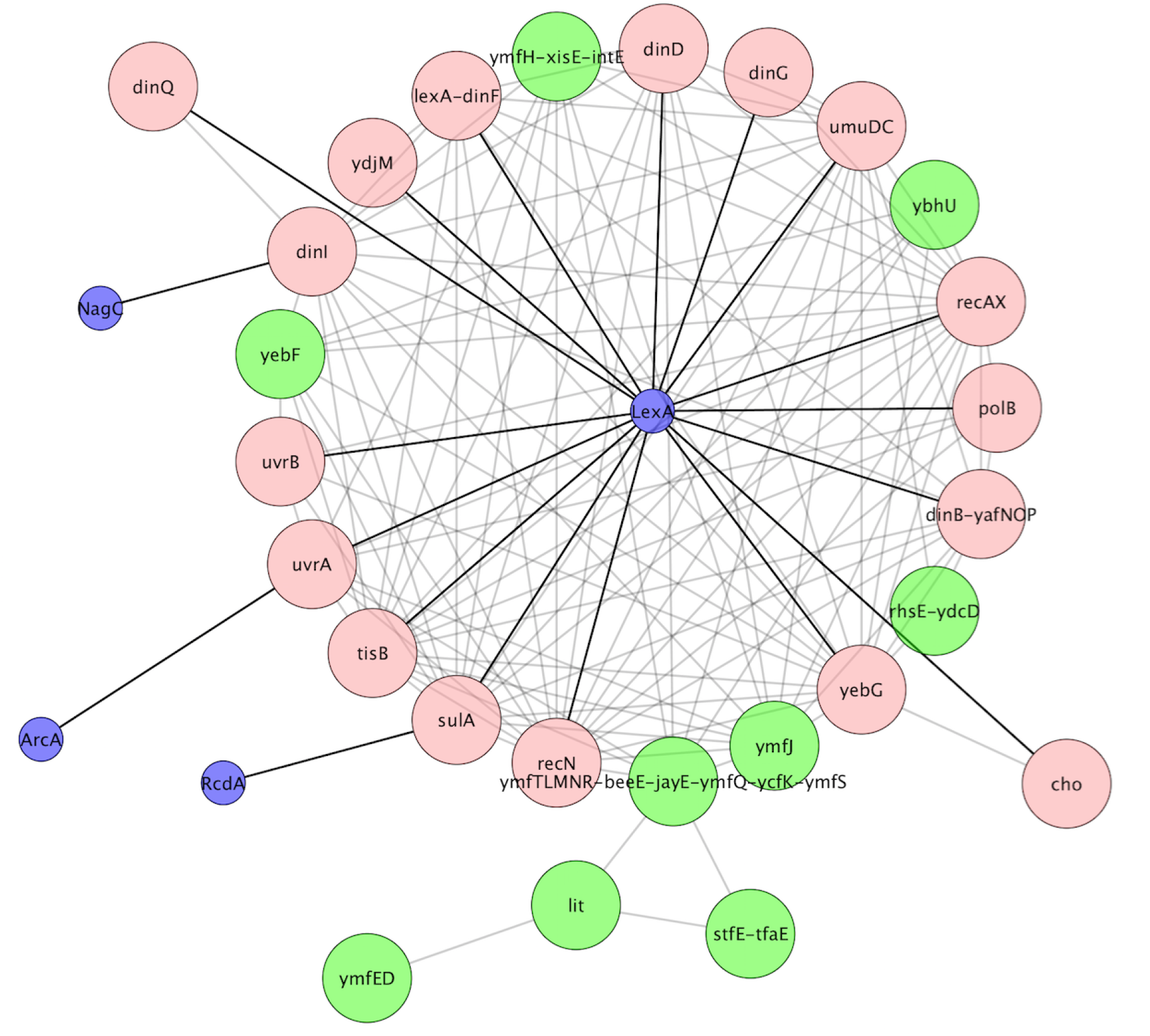
The sub-network of the group ‘LexA’. The group contains 44 genes. Genes belonging to the same operon are collapsed into the same node in the figure. Pink and green nodes are the members in the group. Small blue nodes are TFs which have regulatory connection with the operons (genes) in the group. Links between genes are shown as light gray lines and the regulatory relationship from RegulonDB is drawn as black lines.

**Figure 7:**
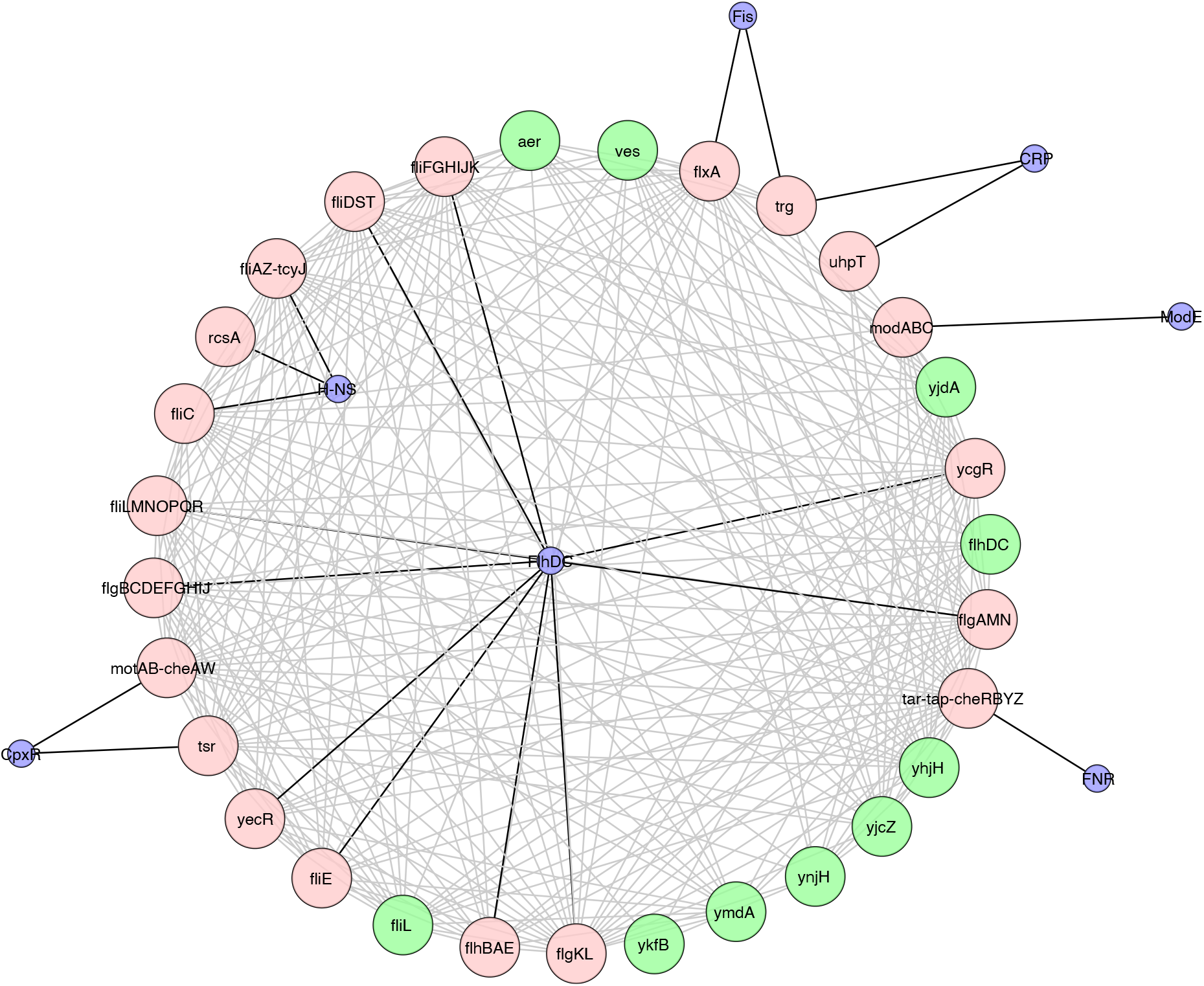
The sub-network of the group ‘Mobility’. The group contains 67 genes. Genes belonging to the same operon are collapsed into the same node in the figure. Pink and green nodes are the members in the group. Small blue nodes are TFs which have regulatory connection with the operons (genes) in the group. Links between genes are shown as light gray lines and the regulatory relationship from RegulonDB is drawn as black lines.

## 4. Conclusions

In this study, we performed clustering analysis on weighted gene regulatory network of *E. coli* and used the three-body correlations instead of ordinary two-body correlations between a pair of genes to highlight certain important gene clusters. This focused analysis reduces noise and allows us to clearly observe hidden patterns in the inferred regulatory network. Using this approach we have identified 25 communities of genes that are relevant from a biological viewpoint. We notice that these communities detected by our approach differ from communities reported in [47] with a computed Normalized Mutual Information (NMI) score of 0.56, considering only the genes present in both partitioning. We posit that this approach of using the weighted links with a threshold and that of considering one gene as the objective gene to infer the gene relationships and groups can potentially be applied to a broader class of clustering problems and can reveal more about the community structure which is otherwise difficult to observe using binary network analysis and two-body correlations. Although some network inference methods produce directed networks describing influence or direct interaction of one gene on another [48], these methods require dynamical expression data.

The inferred gene relatedness networks are far from being random structures, and display non-trivial structural properties. Studies based on gene expression data have shown that the inferred networks exhibit hierarchical [21, 49], scale-free [50, 51] and small world properties [51]. While the information at the system level is available, relatively less is understood about organization of genes in small functional units. This is due to the fact that clustering algorithms and network inference methods applied to gene expression data often have some limitations. Co-expression of genes does not always imply co-regulation, and conversely, lack of co-expression between mRNA expression profiles does not always imply lack of co-regulation [52]. Even the best performing network inference methods can only recover a modest portion of experimentally determined regulatory interactions [12]. Moreover, considerable variability exists in the community detection process due to the approximate stochastic essence of current computational algorithms. Apart from the limitations of methods, we can also ask if the type of connection between genes has an impact. For instance, can genes connected via transcriptional regulation be clustered while genes connected via post-transcriptional regulation can not? In this paper we investigate this question using a robust community detection approach that takes into consideration three-body correlations among genes. We highlight a few test case examples of gene communities that focus on the global regulators *LexA*, *CRP*, *CsrA*, *Fur*, *FlhDG* etc. and their target genes in *E. coli*.

## Acknowledgments

This work was supported by the NSF through grants DMR-1507371 and IOS-1546858. Some of the computations in this work were done on the uHPC cluster at the University of Houston, acquired through NFS Award Number 1531814. We thank Madeleine Opitz for fruitful discussions.

## Supplementary Material

File: S1. robust-gene-communities-25.xlsx

File: S2. go-term-enrichment-communities.xlsx

